# Natural protection from type 1 diabetes in Non Obese Diabetic (NOD) mice is characterised by a unique pancreatic islet phenotype

**DOI:** 10.1101/2021.01.10.426116

**Authors:** Joanne Boldison, Terri C. Thayer, Joanne Davies, F. Susan Wong

## Abstract

The non-obese diabetic (NOD) mouse develops spontaneous type 1 diabetes, with some features of disease that are very similar to the human disease. However, a proportion of NOD mice are naturally-protected from developing diabetes, and currently studies characterising this cohort are very limited. Here, using both immunofluorescence and multi-parameter flow cytometry we focus on the pancreatic islet morphology and immune infiltrate observed in naturally-protected NOD mice. We show that naturally-protected NOD mice are characterised by an increased frequency of insulin-containing, smaller sized, pancreatic islets. Although mice remain diabetes free, florid immune infiltrate remains. However, this immune infiltrate is skewed towards a regulatory phenotype in both T and B-cell compartments. Pancreatic islets have an increased frequency of IL-10 producing B cells and associated cell surface markers. Resident memory CD69^+^CD8^+^ T cells show a significant shift towards reduced CD103 expression, while CD4^+^ T cells have increased FoxP3^+^CTLA4^+^ expression. These data indicate that naturally-protected NOD mice have a unique islet signature and provide new insight into regulatory mechanisms within pancreatic islets.

## Introduction

Type 1 diabetes is an organ-specific autoimmune disease characterized by immune-mediated beta-cell destruction in pancreatic islets, which results in deficient insulin production. Similar to humans, Non Obese Diabetic (NOD) mice develop spontaneous type 1 diabetes. However, in NOD mouse colonies worldwide, approximately 20% (or more) of NOD mice remain normoglycemic and ‘protected’ from diabetes, despite their genetic predisposition (1). Few studies have been done to discover the mechanism of this natural protection. Recently, we have dissected the B-cell functionality in naturally-protected NOD mice, highlighting an increased IL-10-producing B-cell frequency and enhanced response to dendritic cells, compared to NOD mice that have developed diabetes (2). Furthermore, it has been suggested that B cells, specifically anergic CD40^+^IL-10-producing B cells, found in the pancreatic islets of long term normoglycemic mice (protected) (3), may confer this natural protection. Currently, the phenotype and function of CD4^+^ and CD8^+^ T cell populations in naturally-protected NOD mice is unexplored.

Pancreatic islets have a dynamic tissue microenvironment, in which immune cells communicate to drive beta cell destruction. This is complicated by cellular and kinetic heterogeneity in both mouse and human pancreatic islets, including the rate of beta cell destruction. The aim of this study was to investigate the characteristics and heterogeneity of the islets in naturally-protected NOD mice, including the immune infiltrate. This knowledge may provide insight into disease heterogeneity in humans, as not all at-risk individuals develop type 1 diabetes.

## Research Design and Methods

### Mice

Female NOD/Caj mice, originally from Yale University, were bred in-house at Cardiff University. All mice received water, irradiated food *ad libitum* and were housed in specific pathogen-free isolators or scantainers, with a 12h dark/light cycle, at Cardiff University. All animal experiments were approved by Cardiff University ethical review process and conducted under United Kingdom Home Office license in accordance with United Kingdom Animals (Scientific Procedures) Act, 1986 and associated guidelines. Diabetes conversion rates for Cardiff University can be found in (4) with approximately 80% incidence in females by 30 weeks, with a median incidence at 19 weeks old. NOD female incidence at other institutions or companies can vary; e.g. at Jackson laboratories incidence is approximately 90% incidence by 30 weeks of age, with a median female onset at 18 weeks.

### Diabetes Incidence

Mice were monitored weekly for glycosuria (Bayer Diastix) from 12 weeks of age. Following 2 positive glycosuria measurements, blood glucose levels were tested and if greater than 13.9mmol/L, mice were diagnosed as diabetic. NOD mice that were 35 weeks of age or older and had never tested positive for glycosuria were considered to be protected from diabetes, as the incidence of diabetes after this age is very low.

### Islet preparation

Pancreata were inflated with collagenase P solution (1.1mg/ml) (Roche, Welwyn Garden City, UK) in HBSS (with Ca^2+^ and Mg^2+^) via the common bile duct, followed by collagenase digestion with shaking at 37°C for 10min. Islets were isolated by Histopaque density centrifugation (Sigma-Aldrich, Dorset, UK), and hand-picked under a dissecting microscope. For flow cytometric analysis islets were then trypsinized to generate a single cell suspension. Islet cells were rested at 37°C 5% CO2 in IMDM (supplemented with 5% fetal bovine serum (FBS), 2mM L-glutamine, 100U/mL Penicillin, 100μg/mL Streptomycin, and 50μM β-2-mercaptoethanol) overnight, before multiparameter staining.

### Fluorescence immunohistochemistry

Pancreatic tissues were frozen in optimal cutting temperature medium and sectioned at 7µm thickness. For wholemounts, pancreatic islets were fixed overnight at 4°C in 1% paraformaldehyde (PFA). For pancreatic sections, sections were fixed in 1% PFA for 1hr at RT. Following fixation, tissue was permeabilised with 0.2% triton-x100 and blocked with 5% FBS before the addition of a rat anti-mouse CD45 (Biolegend) and a biotinylated anti-insulin (Abcam; clone D6C4,) antibody mix. Secondary labelling was performed with both AlexaFluor 633-conjugated goat anti-rat antibody (Invitrogen) and a Streptavidin-conjugated AlexaFluor 488 antibody (Invitrogen) and mounted with VECTASHIELD® mounting medium, with DAPI (Vector Laboratories). Islet wholemounts were centrifuged at 300g for 3mins before being resuspended in mounting medium, with DAPI, before mounting to the slide. All sections and wholemounts were imaged on Leica SP5confocal microscope.

### Image J analysis

All analysis was performed using Fiji (Image J) (5). Islet area, perimeter, circularity, CD45 and insulin intensity was measured by using a region of interest (ROI) on individual channels using Fiji’s measurement tool. Islets with insulin remaining were considered to be insulin containing islets (ICI) when 3 or more insulin^+^ beta cells were present.

### Flow Cytometry

Cells were incubated with TruStain (anti-mouse CD16/32 [Biolegend, London, UK]) for 10min at 4°C, followed by fluorochrome-conjugated mAbs against cell surface markers for 30min at 4°C. B cell phenotyping multi-parameter flow cytometry was carried out using mAbs: CD3 (145-2C11); B220 (RA3-6B2); CD138 (281-2); CD86 (PO3); CD80 (16-10A1); CD11c (N418); CD11b (M1/70); CD19 (6D5); CD44 (IM7); BAFFR (7H22-E16); MHC II (10-3-6); Ki67 (11F6), all purchased from Biolegend. IL-10 (JES5-16E3); IgD (11-26c.2a); CD40 (3/23) purchased from BD Biosciences. Galectin-1 antibody was purchased from RD systems. T cell phenotyping multi-parameter flow cytometry was carried out using mAbs: CD3 (145-2C11); CD8 (53-6.7); CD4 (GK1.5); CD103 (2E7); CD69 (H1-2F3); PD-1 (29F.1A12); IFNγ (XMG1.2); CTLA4 (UC10-4B9); FoxP3 (MF-14), all purchased from Biolegend. CD25 (PC61); IL-10 (JES5-16E3) were purchased from BD Biosciences. Dead cells were excluded from analysis by Live/Dead exclusion dye (Invitrogen, MA, USA). IFNγ; IL-10; CD107a and Galectin-1 were detected by intracellular cytokine staining after 3hrs of stimulation with PMA (50ng/ml), ionomycin (500ng/ml) and monensin (3µg/ml) [all from Sigma-Aldrich]. After extracellular staining, cells were fixed using fixation/permeabilization kit according to the manufacturer’s instructions (BD Biosciences) and subsequently stained for mAb against intracellular cytokines or appropriate isotype controls. For FoxP3, CTLA4 and Ki67 staining cells were fixed/permeabilized using eBioscience nuclear transcription kit. Cell suspensions were acquired on an LSRFortessa (FACSDIVA software, BD Biosciences), and analysed using Flowjo software, version 10.1 (Tree Star, Oregon, USA).

### Statistical analysis

Statistical analyses were performed using GraphPad Prism (GraphPad Software, San Diego, CA). Comparison between groups was determined by Mann-Whitney U test or Kolmogorov-Smirnov test. For correlations, Pearson correlation co-efficient was calculated. Data were considered significant at p<0.05.

## Data Availability

The datasets generated or analysed during the current study are available on reasonable request.

## Results

### Increased frequency of insulin-containing small pancreatic islets in naturally-protected NOD mice

To investigate the features of the pancreatic islets in naturally-protected NOD mice (not diabetic by 30 weeks of age and hereafter referred to as protected), we used immunofluorescence histochemistry, including both pancreatic islet wholemounts and sections. First, we analysed size, by area measured in pixels/µm, of pancreatic islets in wholemounts, from protected NOD mice (Fig. 1a, b). Representative images (Fig. 1a) and summary graph (Fig. 1b) demonstrated a range of sizes in the remaining islets in protected NOD mice. Next, we determined the size, by area, of pancreatic islet sections, from both protected and diabetic NOD mice (Fig. 1c, d). Smaller pancreatic islets were significantly more frequent in NOD mice that were protected from diabetes, compared to mice that had developed diabetes (Fig. 1d) (p<0.001). To analyse these islet data further, we used a frequency distribution graph to show the relative contribution, in percentage, of each islet to set ‘bins’ according to islet area. Islet size distribution analysis (in percentage) showed that the relative frequency of islets, with an area smaller than 50,000 pixels/µm, was greater in protected (85%) compared with diabetic (51%) NOD mice (Fig. 1e). Further analysis in both protected and diabetic NOD mice revealed that insulin-containing islets (ICI) with detectable insulin-positive beta cells were significantly smaller in size and more frequent in the protected NOD mouse pancreata compared to diabetic NOD mice (p<0.01) (Fig. 1f). Interestingly, in the diabetic NOD mice, the very few ICI detected were larger in area (Fig. 1g). Crucially, a comparison between insulin-containing islets (ICI) and insulin-deficient islets (IDI), in both NOD groups, revealed that ICI were significantly smaller in islet area (Fig. 1h) and were more frequent (Fig. 1i), in comparison to IDI in protected NOD mice. However, this feature was lost in diabetic NOD mice with no significant difference found in islet area (Fig. 1h), or frequency (Fig.1i), in the few ICI identified.

**Figure 1.**
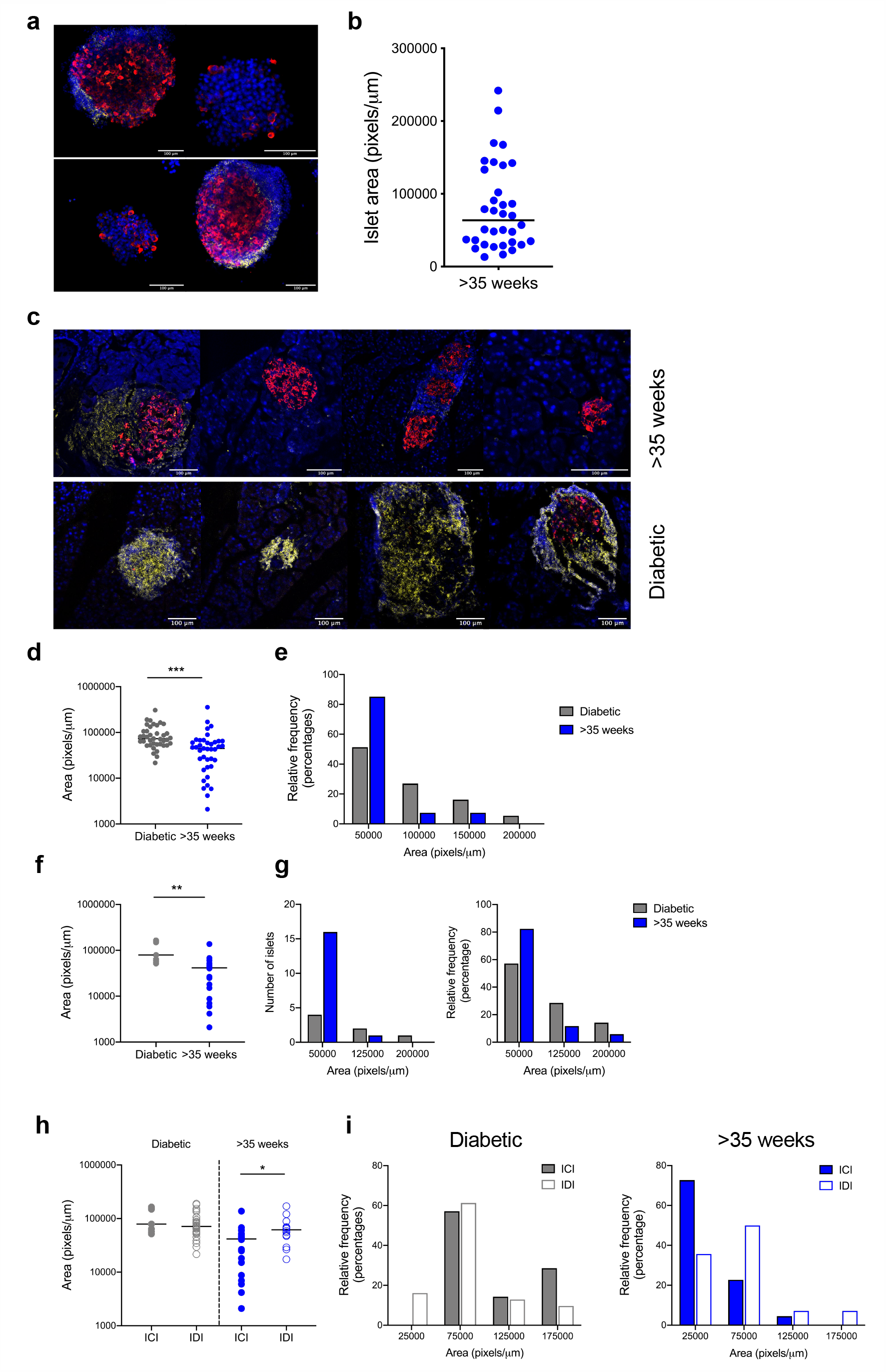
Pancreatic islets from naturally-protected NOD mice are smaller in size. Pancreatic islet wholemounts and sections from naturally-protected NOD mice (>35 weeks old; blue) and NOD mice that had developed diabetes (grey) were analysed by immunofluorescence staining for CD45 (yellow), insulin (red) and DAPI (blue). (a) Representative images of wholemounts from naturally protected >35-week-old mice; (b) graph showing distribution of individual islet area in >35-week-old mice; (c) representative images from pancreatic sections from >35-week-old mice and mice that have developed diabetes; (d) Summary graphs of islet area from individual islets in >35-week-old and diabetic NOD mice (left) and (e) the relative frequency distribution in percentage (right) of pancreatic islets; (f) summary graphs for insulin containing islets (ICI) in both >35-week-old and diabetic NOD mice showing area, (g) frequency distribution in number of islets (left), relative frequency distribution in percentage (right); (h) summary graphs for insulin-containing islets (ICI) and insulin-deficient islets (IDI) from >35-week-old and diabetic NOD mice showing islet area (left); (i) relative frequency distribution of percentage of these islets in mice diabetic NOD mice (left), and relative frequency distribution of percentage of these islets in >35-week-old NOD (right). Data shown are from at least 2 independent experiments (protected n=3; diabetic n=3). *<0.05; **<0.01; ***<0.001; Kolmogorov-Smirnov test.

Alongside islet area, islet perimeter was also analysed and the features observed in protected NOD mice were further confirmed (SFig. 1a-c). In addition to islet size, we investigated whether islet circularity was different in protected NOD mice, compared to mice that have developed diabetes; however no difference was observed in total islets analysed (SFig. 1d) or between ICI and IDI in either the protected or diabetic NOD mouse cohort (SFig. 1e).

### Morphological characterisation of pre-diabetic NOD pancreatic islets

Since protected mice had an increased frequency of smaller islets than diabetic mice, we determined if smaller islets were less affected by immune cell infiltrate when pancreatic insulitis was established and diabetes begins to manifest (mice aged 12-18 weeks old). Wholemount pancreatic islets were stained with insulin and CD45 (a marker of immune cells) to identify islets undergoing attack, and the size was measured by area. Figure 2a demonstrates heterogeneity of pancreatic islets, shown by representative pictures from two different NOD mice. Z-stacks taken from two individual islets are also shown in SFig. 2a. As expected, size, shape and quantity of the CD45 immune infiltrate were variable in each individual islet. However, we observed a clear pattern between size and quantity of CD45 immune infiltrate. We confirmed this with correlation analysis, comparing islet area (Fig. 2b) and islet perimeter (SFig. 2b), with fluorescence intensity of CD45 (left) and insulin (right). A significant positive correlation was observed (p<0.001) between islet area and CD45 immune infiltrate, alongside a significant negative correlation (p<0.01) between islet area and insulin^+^ beta cells.

**Figure 2.**
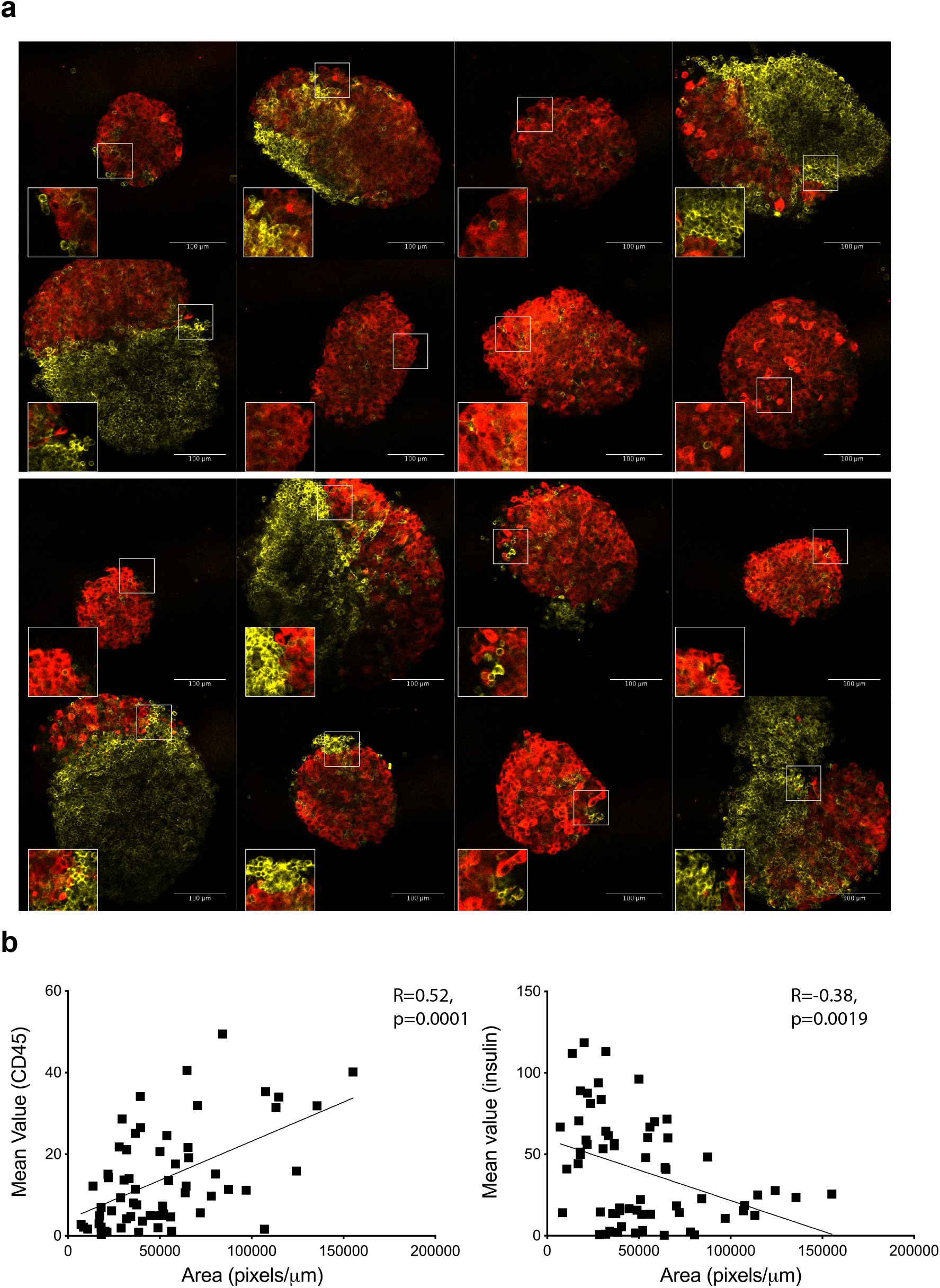
Morphological characterisation of pre-diabetic NOD pancreatic islets. Pancreatic islet wholemounts from NOD mice aged 12-18 weeks were analysed by immunofluorescence staining for the immune cell marker CD45 (yellow) and beta cell marker insulin (red). (a) Representative images from 2 (top panel, bottom panel) individual NOD mice, 8 pancreatic islets shown for each individual NOD mouse. (b) Correlative graphs for both CD45 and insulin expression against islet area. Data are from 4 independent experiments, n=8.

### Regulatory B cells are increased in the islets of naturally-protected NOD mice

As the pancreatic islets of naturally-protected NOD mice have considerable immune infiltration, although remaining normoglycemic, we analysed the B cell infiltrate to investigate the frequency of regulatory cells by flow cytometry (Figure 3). Firstly, we compared the percentage of total B cells in groups of pooled protected NOD mice (>35 weeks), compared to groups of pooled younger pre-diabetic NOD mice (6-8 weeks old) (Fig. 3a, b) and observed a significant increase in CD19^+^ B cells (p<0.001). Secondly, complementing our previous observation that naturally-protected NOD mice have increased splenic IL-10-producing B cells (2), we showed a significant increase in pancreatic islet IL-10-producing B cells (Fig. 3c, SFig. 3a). Furthermore, we also demonstrated a population of Galectin-1^+^ B-cells (SFig. 3b), which encompassed the majority of the IL-10-producing B cells and were increased in >35-week-old mice (p=0.06) (Fig. 3d, SFig. 3c). Interestingly, we also observed a significant increase in CD40^+^ (Fig. 3e) and CD80^+^ expressing B cells (Fig. 3f) (for representative gating see SFig. 3d), both markers associated with regulatory B cell function (6; 7). It should be noted that no significant differences were observed in B cell expression of MHC II, CD86 or BAFFR (SFig. 3e,f).

**Figure 3.**
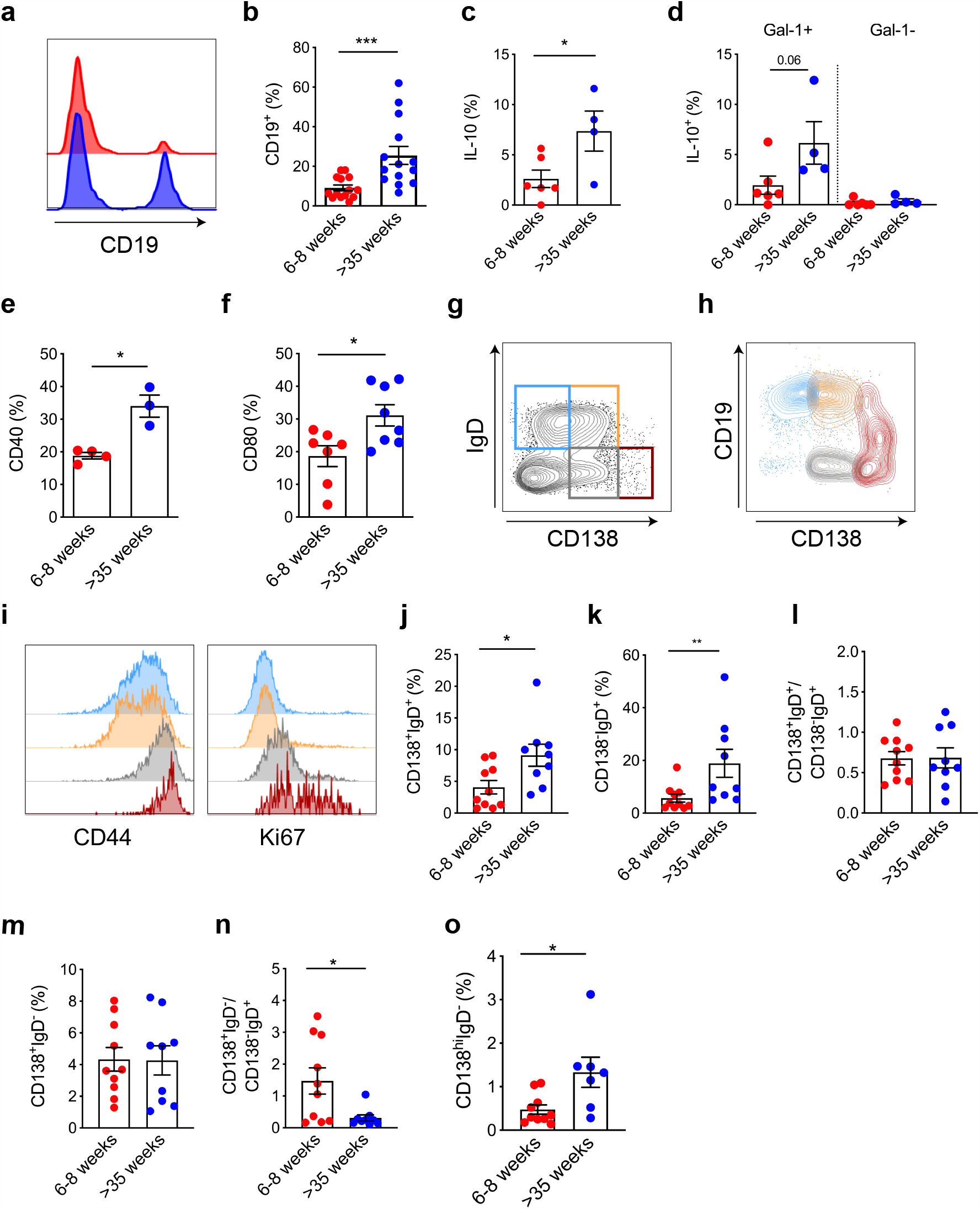
Naturally-protected NOD mice have increased regulatory B cells in the pancreatic islet infiltrate. Pancreata from groups (n=2-3) of NOD mice aged 6-8 weeks old (younger prediabetic; red) and >35 weeks old (protected; blue) were taken and pancreatic islets pooled together before flow cytometric analysis. (a) Representative histograms to show CD19^+^ B cell expression; (b) Overall percentages for CD19^+^ B cells; (c-f) Overall percentages of CD19^+^ B cells expressing (c) IL-10, (d) IL-10^+^ in Gal-1^+^ and Gal-1^-^ compartments, (e) CD80 (f) CD40; (g-i) representative flow cytometry plots gated on live, CD3^-^CD11b^-^CD11c^-^ cells for (g) CD138 and IgD populations; CD138^-^IgD^+^ (blue gate); CD138^+^IgD^+^ (orange gate); CD138^+^IgD^-^ (grey gate), CD138^hi^IgD^-^ (red gate); (h) CD19 expression on each of the CD138/IgD populations; (i) histograms showing CD44 (left) and Ki67 (right) on each of the CD138/IgD populations; (j, k); Overall percentages for (j) CD138^+^IgD^+^ cells (k) CD138^-^IgD^+^; (l) Ratio of CD138^+^IgD^+^ to CD138^-^IgD^+^ B cells; (m) Overall percentages of CD138^+^IgD^-^ cells; (n) Ratio of CD138^+^IgD^-^ cells to CD138^-^IgD^+^ B cells (o) Summary graph for percentages of CD138^hi^IgD^-^ plasmablasts in both young NOD and protected >35-week-old NOD mouse groups. Data shown are from at least 4 independent experiments. *<0.05; **<0.01; ***<0.001, Mann-Whitney U test.

Many B cells in pancreatic islets express CD138 (8; 9), a marker that identifies plasmablasts and plasma cells. We investigated our previously described populations identified by CD138 and IgD expression (9), we examined these populations further (Fig. 3g-i), using markers for murine plasmablasts (10; 11). We assessed CD19 (Fig. 3h), CD44 and Ki67 (Fig. 3i) in each CD138^+/-^ population revealing that CD138^hi^IgD^-^ cells (red gate) contained a CD44^hi^Ki67^+^ highly proliferative population which also expressed intermediate levels of CD19 (CD19^int^). With this further analysis, we propose that CD138^hi^IgD^-^ cells (red gate) are a subpopulation of dividing plasmablasts. Few B cells that remained IgD^+^ were proliferating and so it is probable that they represent classical B cells (blue gate, CD138^-^IgD^+^) and an intermediate-stage of plasmablast (orange gate, CD138^+^IgD^+^). CD138^+^IgD^-^ cells (grey gate) were a mixture of both +/-CD19 cells and increased Ki67 expression, compared to CD138^-^IgD^+^ classical B-cells, and are likely to represent both non-dividing intermediate-stage plasmablasts and dividing plasmablasts.

We demonstrated that protected NOD mice displayed significant increases in both CD138^+^IgD^+^ B cells (Fig. 3j), and CD138^-^IgD^+^ B cells (Fig. 3k), compared to young 6-8-week-old mice. This observation indicated that CD138^+^IgD^+^ cells were not selectively recruited over CD138^-^IgD^+^ B-cells (Fig. 3l). However, the frequency of CD138^+^IgD^-^ B-cells in both young and older, protected NOD mouse groups was not altered (Fig. 3m). Thus, the ratio of the CD138^+^IgD^-^ to CD138^-^IgD^+^ B cells was increased in the younger NOD mice (Fig. 3n). This may suggest these B cells arrive early to the pancreatic islets. Analysis of the small population of dividing plasmablasts (CD138^hi^IgD^-^) showed a significant increase in protected NOD mice, compared to younger NOD mice (Fig. 3o).

### Enrichment of CD4^+^FoxP3^+^CTLA4^+^ Tregs in naturally-protected NOD mice

To determine the characteristics of CD4^+^ T cells in the pancreatic islets of NOD mice, that are ‘naturally-protected’ from diabetes, we studied the islet-infiltrating T cells of groups of pooled protected NOD mice by multiparameter flow cytometry, and compared these to islet-infiltrating T cells from groups of pooled mice 6-8 weeks of age (mice with early-stage insulitis). CD4^+^ T cell frequency and expression were increased in protected NOD mice, although non-significant due to the variability in younger NOD mice (Fig. 4a, b). To ascertain if CD4^+^ Tregs contributed to the protection seen in protected NOD mice, we investigated the presence of CD4^+^FoxP3^+^ Tregs in the pancreatic islets and revealed a significant increase in CD4^+^FoxP3^+^ Treg cells in the pancreatic islets of protected NOD mice, compared to 6-8-week-old mice (Fig. 4c, d). Further analysis of the CD4^+^FoxP3^+^ T cells revealed a significant increase in the frequency of CTLA4^+^CD4^+^FoxP3^+^ T cells in protected NOD mice, compared to 6-8-week-old mice (p<0.01) (Fig. 4e, f). No significant differences were observed in the percentage of CD4^+^FoxP3^+^ T cells expressing CD25, PD-1 or CD103. Interestingly, intracellular cytokine analysis of both CD4^+^FoxP3^-/+^ T cells showed a significant increase in IFNγ-producing CD4^+^FoxP3^-^ T cells in protected NOD mice, compared to mice aged 6-8 weeks old (Fig. 4g,h). No differences in IL-10 in FoxP3^+^ CD4^+^ T cells was observed. This increase in IFNγ production from re-stimulated CD4^+^ T cells may reflect enhanced antigen experience.

**Figure 4.**
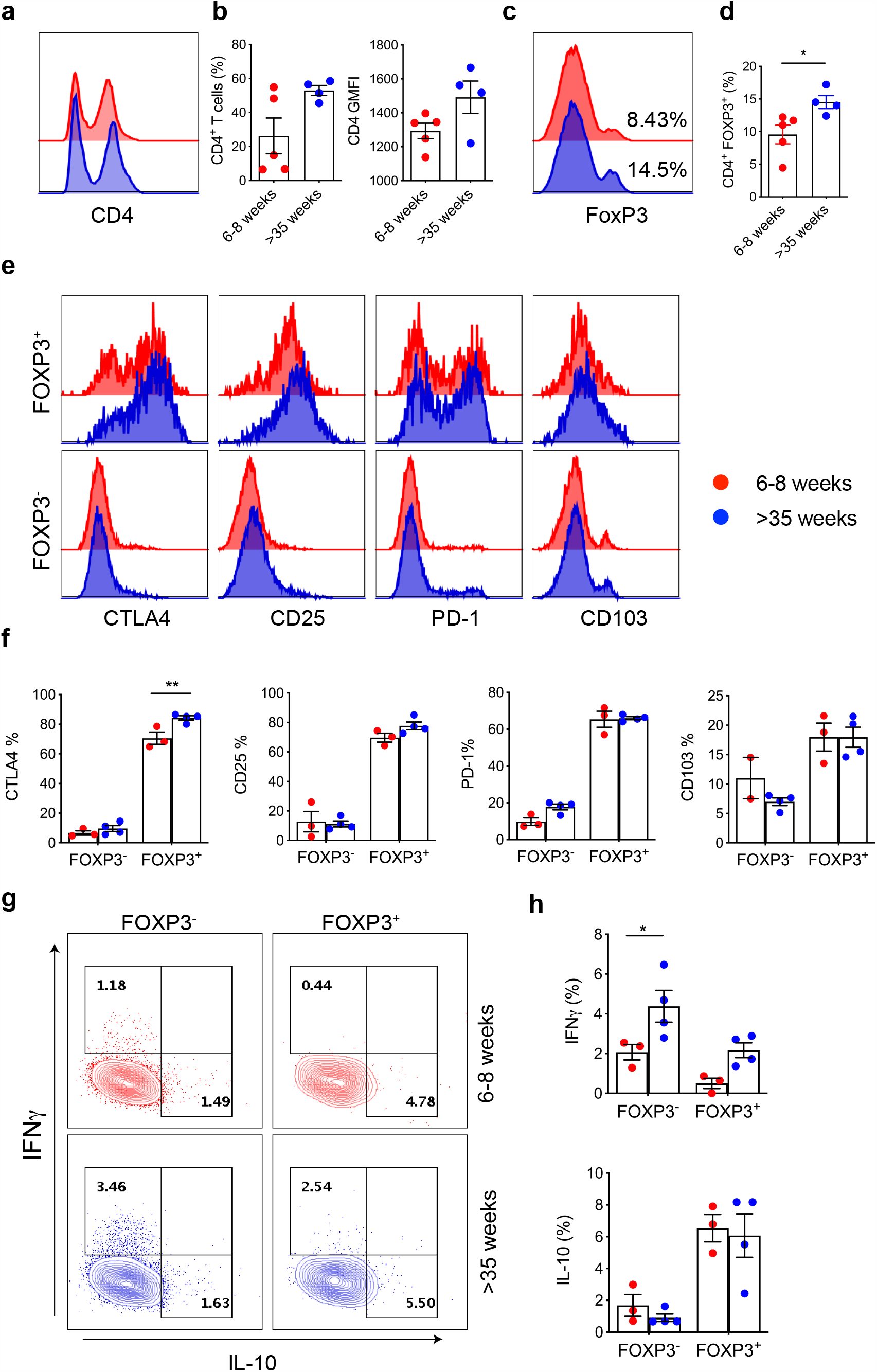
Increased frequency of CD4^+^CTLA4^+^FoxP3^+^ Tregs in naturally-protected NOD mice. Pancreata from groups (n=2-3) of NOD mice aged >35 weeks old (protected; blue) and 6-8 weeks old (younger prediabetic; red) were taken and pancreatic islets pooled together before flow cytometric analysis. (a) representative histogram of CD4 expression; (b) CD4^+^ T cell percentages (left) and CD4 geometric mean fluorescence intensity (GMFI) (right); (c) CD4^+^FoxP3^+^ T cell expression representative histograms and (d) summary graph; (e, f) expression of CD4^+^ Treg markers on CD4^+^FoxP3^+^ and CD4^+^FoxP3^-^ T cells with (e) demonstrating representative histograms for CTLA4, CD25, PD-1 and CD103 and (f) graphs showing summary percentages; (g, h) IFNγ and IL-10 expression in CD4^+^FoxP3^+/-^ subsets, with (g) illustrating representative flow plots and (h) summary graphs. Cells were gated on singlets, live, CD4^+^ cells. Data are from at least 3 independent experiments. *<0.05; **<0.01, Mann-Whitney U test.

### Islet CD8 T_RM_ in naturally-protected NOD mice switch to a CD103^-^ phenotype

Additional analysis of T cells was performed. Similar to CD4^+^ T cells, CD8^+^ T cells were modestly increased in frequency in >35-week-old mice (Fig.5a,b). As CD8^+^ T cells are found from an early age in the pancreatic tissue we investigated markers of tissue residency. CD8^+^ tissue resident memory cells (T_RM_) can be distinguished by the surface markers CD69 and CD103 (12). CD8^+^ T_RM_ populations have now been widely studied in various tissues and play a crucial role in immunosurveillance and protect against secondary viral infections (13). We demonstrated 3 key populations of CD8^+^ T cells in the pancreas of NOD mice: CD103^-^CD69^-^ recirculating cells; CD103^-^CD69^+^ T_RM_ and CD103^+^CD69^+^ T_RM_ (Fig. 5d-g). Protected >35-week-old NOD mice and younger NOD mice had similar frequencies of recirculating CD8^+^CD103^-^CD69^-^ T cells (Fig. 5d). Strikingly, CD8^+^ T_RM_ cells were significantly different between protected NOD mice and mice aged 6-8 weeks old, with a shift towards greater prominence of CD103^-^ T_RM_ cells in protected NOD mice (Fig. 5e, f). Further, both CD8^+^ CD103^-/+^ T_RM_ populations expressed CD107a, IFNγ and PD-1, with the CD103^-^ T_RM_ becoming more activated overall, after stimulation (Fig. 5g). There was no enrichment of PD-1 (a marker for T cell exhaustion (14) in either T_RM_ population (Fig. 5g). Further analysis of the CD107a^+^ cells (a marker for recent degranulation) in the T_RM_ populations, revealed CD107a^+^CD103^+^ T_RM_ cells had fewer IFNγ^+^PD-1^-^ expressing cells than the CD107a^+^CD103^-^ T_RM_ (P<0.05) (Fig. 4i, summary graph), but overall a greater proportion of PD-1 expression (Fig. 4h, pie charts). Evaluation of IFNγ and PD-1 sub-populations in the younger as well as protected NOD mice showed a significant increase in IFNγ^+^PD-1^+^ T cells in both CD103^+/-^ T_RM_ populations (Fig. 4j, k). Overall, naturally protected NOD mice show a shift towards a CD8^+^CD103^-^ T_RM_ population, which has a more activated phenotype after stimulation, alongside an increase in IFNγ in the T_RM_ subsets.

**Figure 5.**
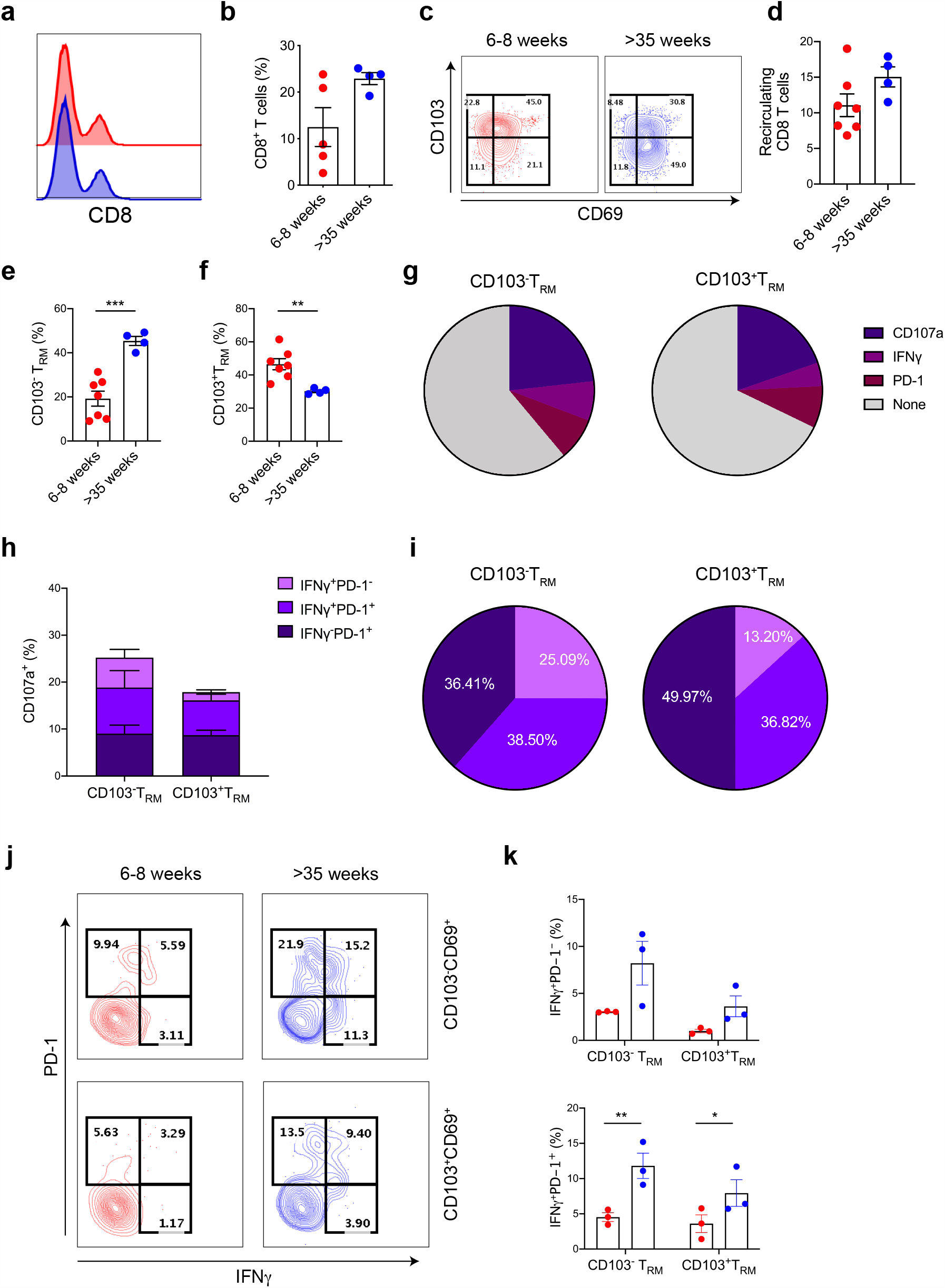
CD8^+^ T_RM_ in naturally-protected NOD mice switch to a CD103^-^ phenotype. Pancreatic islets from groups (n=2-3) of NOD mice aged >35 weeks old (protected; blue) and 6-8 weeks old (younger prediabetic; red) were taken and pancreatic islets pooled together, before flow cytometric analysis. (a) representative flow plots showing CD8^+^ T cells; (b) CD8^+^ T cell percentages; (c) representative flow plots to show CD103^-^CD69^-^ recirculating CD8^+^ T cells and CD103^+/-^CD69^+^ T_RM_ cells (d-g) percentages CD8^+^ T cell populations with (d) demonstrating CD103^-^CD69^-^ recirculating T cells, (e) CD103^-^CD69^+^ T_RM,_ (f) CD103^+^CD69^+^ T_RM_ and (g) pie charts summarising total percentages of CD107a, IFNγ and PD-1 expression on both CD103^+/-^ T_RM_ CD8^+^ subsets; (h, i) CD8^+^CD103^+/-^ T_RM_ subset analysis gated on CD107^+^ cells containing IFNγ^+^PD-1^-^; IFNγ^+^PD-1^+^ and IFNγ^-^PD-1^+^ subsets with (h) showing overall percentages of IFNγ and PD-1 populations and (i) illustrating pie charts showing proportion of CD107a-expressing cells for IFNγ and PD-1 populations (j, k) IFNγ^+^ and PD-1^+^ populations on the CD107a^+^ subset of CD8^+^CD103^-^CD69^+^ T_RM_ (top) and CD8^+^CD103^+^CD69^+^ T_RM_ (bottom) from 6-8-week-old (red) and >35-week-old (blue) NOD mice (j) representative flow cytometry plots (k) summary graphs of percentages. Cells were gated on singlets, live, CD8^+^ cells. Data are from at least 3 independent experiments. *<0.05; **<0.01; ***<0.001, Mann-Whitney U test.

## Discussion

In this report, we demonstrate key characteristics of pancreatic islets in a group of mice that are naturally protected from developing spontaneous diabetes. Firstly, we discovered that smaller islets remain in these protected mice, with a clear correlation between islet size and immune infiltrate. Furthermore, insulin^+^ beta cells are still present in pancreatic islets despite florid immune infiltrate. This immune infiltrate has a high frequency of B and T cells, however the compositional signature was notably different in both immune cell compartments.

For the first time, we show that protected NOD mice have an increased frequency of smaller islets remaining in the pancreas, with insulin-containing islets smaller in size compared to islets deficient in insulin^+^ beta cells. NOD mice that developed diabetes did not display this pattern. Moreover, we show in pre-diabetic NOD mice, larger islets have larger immune infiltrates. Previously, islet size has been shown to decrease as the duration of disease progresses in the NOD mouse (15); however more sophisticated imaging identified that the smaller islets, located peripherally, are destroyed earlier in the disease process (16). However, surprisingly, the CD3^+^ immune infiltrate was not localized to the smaller islets or within a particular islet region (16). In humans, individuals with recent-onset type 1 diabetes have larger islets compared with people with long standing type 1 diabetes (17).

Islets smaller in size than those found in the 12-16 week old NOD mice may have been destroyed; however we noted that islet size and infiltrate were correlated, but curiously the very few remaining islets with insulin^+^ beta cells were not smaller in size. Certainly, no correlation between beta-cell mass and insulitis was observed in human pancreatic sections from donors with type 1 diabetes, but islet size was not addressed (18). This dichotomy requires further study to ascertain why this is the case. An explanation for this correlation between islet size and immune infiltrate could be explained by an increased capillary density (19), providing more immune cell access.

IL-10-producing B cells (B10 cells) diminish the inflammatory response (20). Building on previous work (2), we now show that naturally-protected NOD mice have a regulatory B cell bias in the pancreatic islets. We demonstrate Galectin-1^+^ B cells in pancreatic islets, a marker shown to be necessary for the function of regulatory B cells (21), and production of this protein, from activated B cells, can influence T cell responses, including inducing T cell apoptosis (22). Furthermore, the infiltrating B cells have increased levels of other cell surface markers associated with regulation. Firstly, the CD80 molecule is known to preferentially bind CTLA4 (23), which we also find significantly increased on CD4^+^FoxP3^+^ Tregs in these protected NOD islets. Secondly, B10 cells require CD40 in order to suppress effector T cells and autoimmunity (6). Interestingly, here we describe a significant increase in dividing CD138^hi^CD44^hi^ plasmablasts in protected NOD mice, previously reported to be an IL-10-producing population (10).

We observed that CD138^+^IgD^-^ pancreatic islet B cells are similar in frequency in both younger and protected NOD mice. Early islet B cells have a B1 B cell phenotype (24), B cells that are preferentially located in the peritoneum and pleural cavities, and are players in the initiation of type 1 diabetes (25). CD138^+^IgD^-^ cells identified here are a heterogenous pool of dividing and non-dividing plasmablasts at various intermediate differentiation stages, which consist of CD19^+^ and CD19^-^ cells which lack IgM expression and are more proliferative compared to classical B cells. Crucially, this subset also contains antigen-specific B cells (9). Further work on these defined subsets is required to understand if they are a B1-like-cell that has altered due to the inflamed tissue environment.

Lastly, we identified a significant shift in the CD8^+^ T cell compartment. A CD8^+^CD103^+^T_RM_ phenotype is more prevalent in the younger NOD, whereas a CD8^+^CD103^-^ T_RM_ phenotype dominates in the protected NOD mice, with CD8^+^CD103^-^ T_RM_ cells producing increased IFNγ, when re-stimulated *ex vivo*. CD8^+^ T_RM_ respond rapidly to control local immune responses in the tissue and so are central in tissue immunosurveillance. CD8^+^CD103^-^ T_RM_ have been described in various tissues, with CD8^+^CD103^+^ T_RM_ cells located preferentially in the epidermis while CD8^+^CD103^-^ T_RM_ cells are located in the dermis of human skin (26), with distinct functional roles (26; 27). CD8^+^CD103^+/-^ T_RM_ have been defined as separate populations, with different patterns of recall, molecular signature and migration (27-29). CD8^+^CD103^-^ T_RM_ are more transient in the tissue compared to CD103^+^ counterparts (28), alongside an increased expression of sphingosine-1-phosphate receptor-1 (S1P1R) (27). It remains unclear if the enrichment of the CD8^+^CD103^-^ T_RM_ population in the older protected mice represents a more transient CD8 T_RM_ population, and this would require further interrogation. CD8^+^ T_RM_ in human pancreatic tissue from adults with recent onset type 1 diabetes have been identified, but interestingly, only CD8^+^CD103^+^ T_RM_ cells were detected in pancreatic tissue (30). However, in non-diabetic donors a CD8^+^CD103^-^ T_RM_ phenotype has been observed in approximately 20-30% of resident CD8^+^ T cells in pancreatic islets (31). E-cadherin, the principal ligand for CD103, is expressed by pancreatic islets (32) and this interaction can result in the release of cytokines and lytic granules from CD8^+^ T cells, therefore controlling cytotoxic CD8^+^T cell responses (33; 34). CD103 is also required for the efficient destruction of pancreatic islet allografts (35). A shift towards CD8^+^ T cells lacking CD103 in naturally-protected NOD mice may result in reduced targeted cell death. However, this CD103^-^ T_RM_ shift may also be as a result from the loss of the ligand E-cadherin due to substantial loss of pancreatic islets.

In conclusion, naturally-protected NOD mice have a unique pancreatic signature, with remaining islets that contain smaller insulin-producing beta cells and an immune infiltrate (both T and B cells) shifted towards a regulatory phenotype. These results are important to understand the balance between a destroyed islet and an islet that remains, even partially, intact. Limitations in this study reflect the difficulty of studying both heterogenous islets and immune cell subsets, especially with limited cell numbers. Furthermore, protected NOD mice represent a pool of mice that cannot be detected early in the disease process and so comparisons with mice that are younger, including mice that have established insulitis, is challenging. These younger mice would encompass both mice that will develop diabetes and be naturally protected, and we cannot, as yet, predict which mice will develop diabetes and distinguish them from mice that will be spared. Nevertheless, these observations highlight the need for further investigation into the dynamic process of beta-cell destruction.

## Acknowledgements

This work was funded by the Medical Research Council (UK) MR/K021141/1 to FSW. FSW is the guarantor of this work and, as such, had full access to all the data in the study and takes responsibility for the integrity of the data and the accuracy of the data analysis. No conflict of interest exists for F.S. Wong, J. Boldison, T. Thayer or J.Davies.

## Author contributions

J. Boldison and F.S. Wong designed the experiments and wrote the manuscript. J. Boldison performed the experiments and analyzed the data. J. Davies and T. Thayer contributed to experimental procedures. All authors reviewed the manuscript.

