## Supplementary figures and images for "Natural protection from type 1 diabetes in Non Obese Diabetic (NOD) mice is characterised by a unique pancreatic islet phenotype"

### Supplementary data

##### Supplementary Figure 1

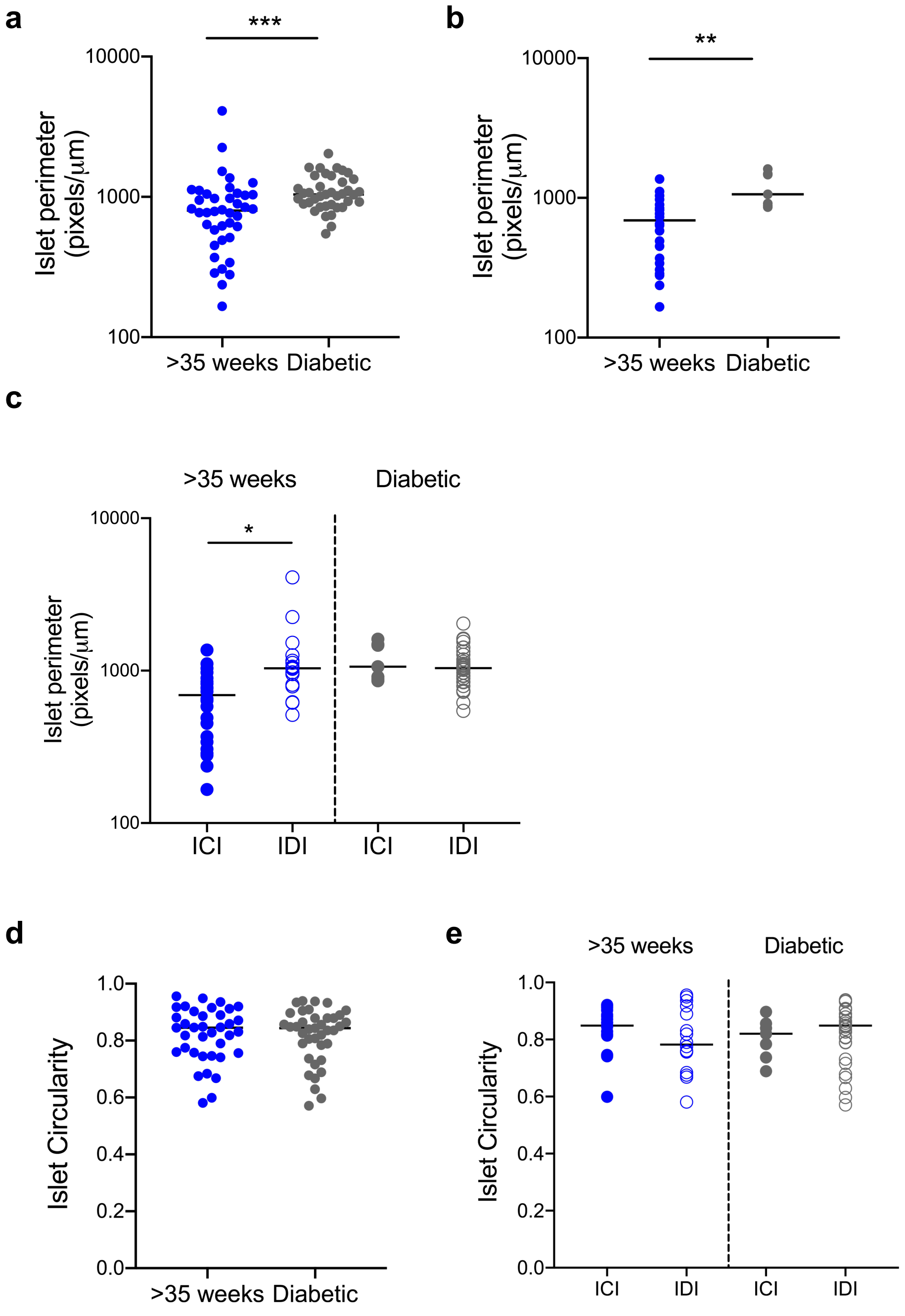

#### Supplementary Figure 2

**a**

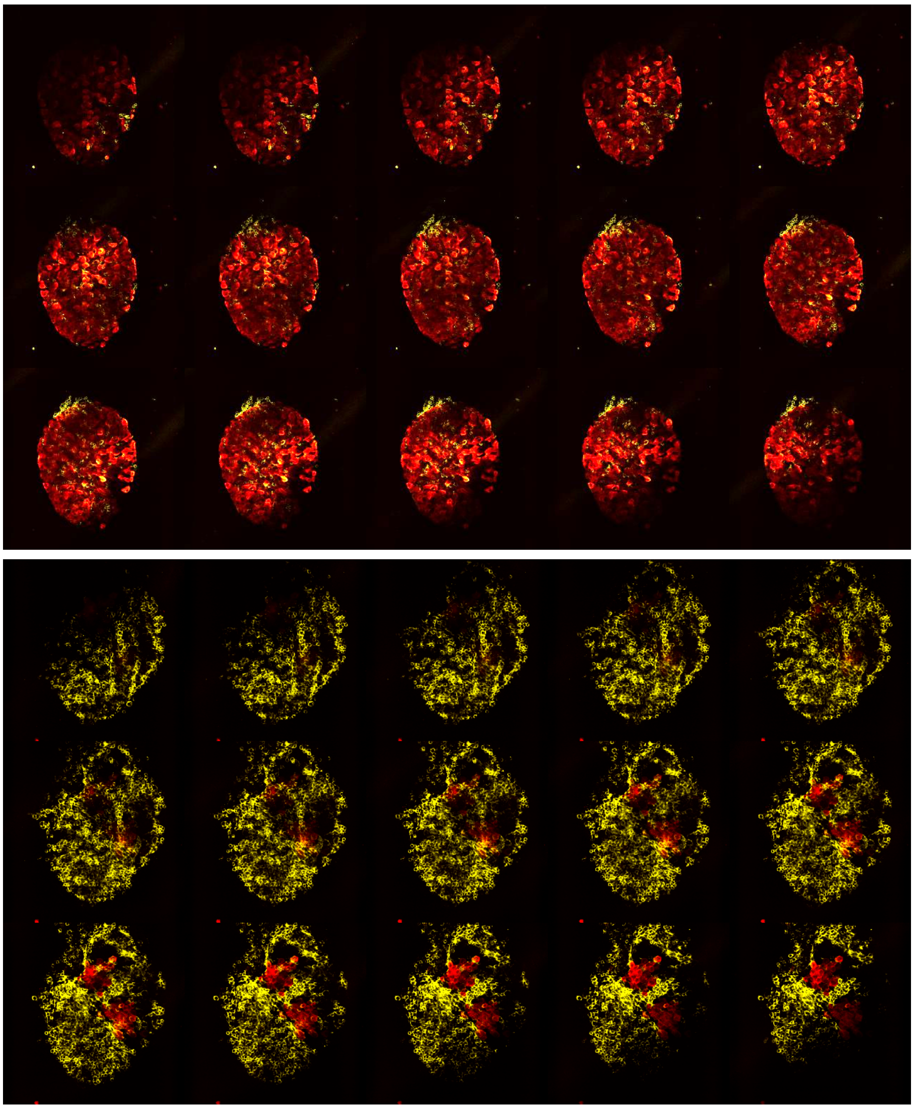

**b**

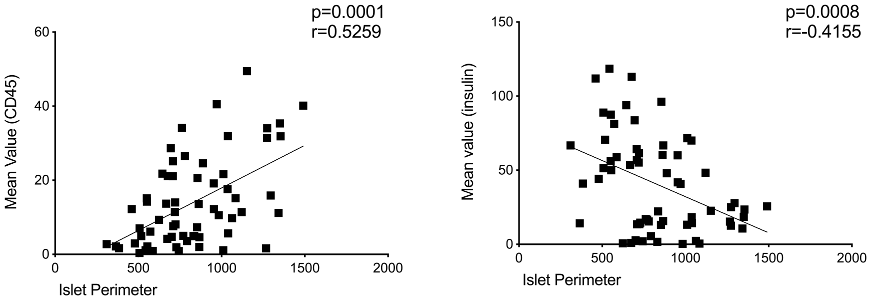

##### Supplementary Figure 3

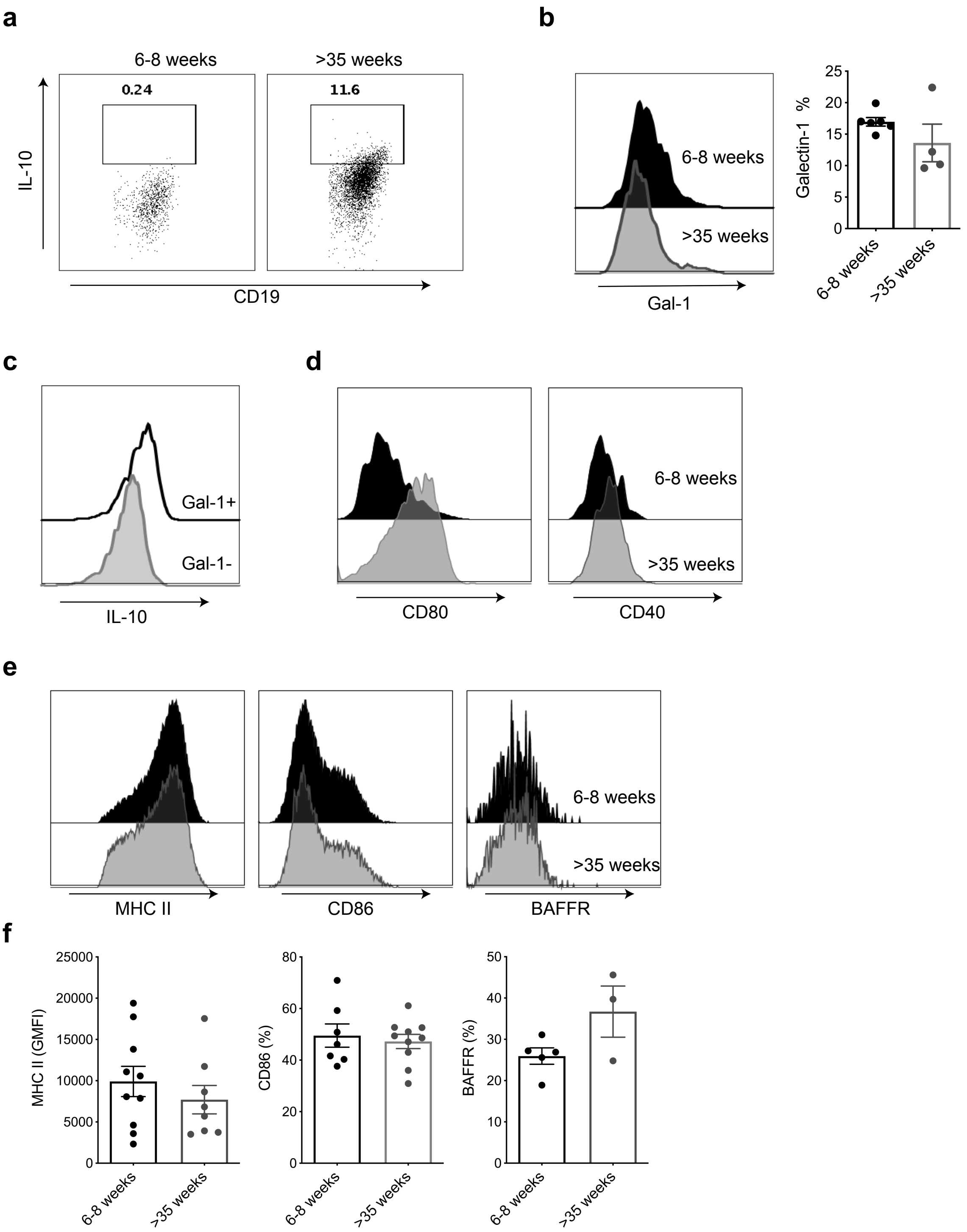
